# Uncovering and exploiting the return of voluntary motor programs after paralysis using a bi-cortical neuroprosthesis

**DOI:** 10.1101/2023.03.01.530610

**Authors:** Maude Duguay, Marco Bonizzato, Hugo Delivet-Mongrain, Nicolas Fortier-Lebel, Marina Martinez

## Abstract

Rehabilitative and neuroprosthetic approaches after spinal cord injury (SCI) aim to reestablish voluntary control of movement. Promoting recovery requires a mechanistic understanding of the return of volition over action, but the relationship between re-emerging cortical commands and the return of locomotion is not well established. We introduced a neuroprosthesis delivering targeted bi-cortical stimulation in a clinically relevant contusive SCI model. In healthy and SCI cats, we controlled hindlimb locomotor output by tuning stimulation timing, duration, amplitude, and site. In intact cats, we unveiled a large repertoire of motor programs. After SCI, the evoked hindlimb lifts were highly stereotyped, yet effective in modulating gait and alleviating bilateral foot drag. Results suggest that the neural substrate underpinning motor recovery had traded-off selectivity for efficacy. Longitudinal tests revealed that the return of locomotion after SCI was time-locked with recovery of the descending drive, which advocates for rehabilitation interventions directed at the cortical target.

**Highlights:**

- A bilateral cortical implant allowed for the delivery of alternate bilateral stimulation coherently with locomotion, which modulated gait trajectories.
- We analyzed the effects of stimulation parameters - timing, duration, amplitude, and site of stimulation - to maximize the improvement of locomotor output after paralysis.
- A varied repertoire of motor programs evoked in intact cats was reduced to one stereotyped response after spinal cord injury (SCI) consisting in flexion modulation that efficiently alleviated hindlimb dragging.
- After SCI, the return of cortical gait control emerged in synchrony with locomotor recovery.

## 1. Introduction

Vertebrate locomotion is controlled by the following mechanisms: (1) integrated spinal central pattern generators (CPGs) that establish motor rhythms (Grillner and Zangger 1979, Barbeau and Rossignol 1987, Kiehn 2006, D’Angelo, Thibaudier et al. 2014, Frigon, Thibaudier et al. 2015, Frigon, Desrochers et al. 2017); (2) sensory feedback, which is a reaction to environmental perturbations (Zehr and Stein 1999, Prochazka, Gritsenko et al. 2002, Rossignol, Dubuc et al. 2006); and (3) supraspinal commands, which are involved in locomotion initiation and modulation as well as balance and voluntary control of movement (Drew, Jiang et al. 2002, Jordan, Liu et al. 2008, Fortier-Lebel, Nakajima et al. 2021).

Among supraspinal contributors, the motor cortex is a key player in the execution and voluntary control of hindlimb movements, which has been dissected through various lesions (Jiang and Drew 1996, Metz, Dietz et al. 1998) and recording studies (Sahrmann, Clare et al. 1984, Widajewicz, Kably et al. 1994, DiGiovanna, Dominici et al. 2016). In anesthetized preparations, intracortical microstimulation (ICMS) of the motor cortex evokes simple hindlimb movements (Neafsey, Bold et al. 1986, Hatanaka, Nambu et al. 2001, Seong, Cho et al. 2014, Brown and Martinez 2018). In healthy conscious rats and cats, the application of ICMS during locomotion produces phase-dependent contralateral movements through direct and indirect projections to the spinal cord (Bretzner and Drew 2005, Bonizzato and Martinez 2021, Fortier-Lebel, Nakajima et al. 2021).

Disruption of descending projections due to spinal cord injury (SCI) induces paralysis and loss of locomotor control, whose recovery is a high priority for individuals with lived experience (Ditunno, Patrick et al. 2008). SCI in humans primarily results from contusions and induces variable spinal tract damage that impairs residual pathway functionality (Ahuja, Wilson et al. 2017, RHI 2018). Since 70% of contusive SCIs are incomplete (RHI 2018, N.S.C.I.S.C 2019), supraspinal centers often retain connections with spinal circuits. Although the neuroplasticity of lumbar spinal networks may support the return of locomotor rhythms (Barbeau and Rossignol 1987), sufficient sparing of descending tracts is required for functional return of volitional walking (Delivet-Mongrain, Dea et al. 2020). Motor cortex plasticity plays a prominent role in the recovery of voluntary leg control (Topka, Cohen et al. 1991, Brown and Martinez 2018, Urbin, Royston et al. 2019, Bonizzato and Martinez 2021, Brown and Martinez 2021), although the cortical contribution to recovery from severe contusion in clinically relevant animal models remains poorly understood. In a rat model of thoracic hemisection with paralysis of one leg, we showed that ICMS delivered in phase coherence with ongoing locomotion immediately restored locomotion (Bonizzato and Martinez 2021, Martinez 2022). However, the utility of neuromodulation strategies targeting the motor cortex to immediately restore bilateral control of locomotion after severe contusions is unknown.

To address this knowledge gap, we designed a neuroprosthetic platform whereby ICMS was delivered alternately to the left and right motor cortex during ongoing locomotion. In healthy cats, we extensively characterized the impact of varying stimulation parameters (timing, duration, amplitude, and site of stimulation) on locomotor output. When delivered at the initiation of the leg flexion phase of locomotion (in “phase coherence” with locomotion), ICMS evoked a variety of motor synergies. After contusive SCI that initially paralyzed both legs, bi-cortical stimulation immediately reversed bilateral foot drag and flexion deficits. We also demonstrated that the return of cortically evoked movements simultaneously occurred with the recovery of voluntary motor control. Our data provide a proof-of-concept methodology to independently modulate bilateral limb trajectories, scrutinizing the level, variety, and timeline of cortical control of movement expression. Overall, this study provides the building blocks for stimulation protocols to control and improve gait following severe SCI.

## 2. Materials and methods

### 2.1. Study design

The objective of this research was to determine the immediate effects of ICMS on locomotor output under intact conditions and after a spinal contusion. Three female tabby cats (one-year-old; MBR Waverly LLC, USA) weighing 3-3.5 Kg were first selected for their ability to walk regularly and continuously for several minutes (10–15 min) on a motor-driven treadmill at the speed of 0.4 m/s. The cats were housed together in a 36 m^2^ room with a 12 h light/dark cycle and had free access to food and water. Cats were implanted with electrode arrays within the hindlimb representation of both motor cortices. In addition, intra-muscular electrodes were implanted in the flexor and extensor muscles in both hindlimbs. The immediate effects of uni-cortical and bi-cortical microstimulation on treadmill locomotion was tested using different stimulation parameters (timing, burst duration, amplitude, and site). Cats then received a contusive SCI at spinal level T10. We next evaluated the immediate effects of cortical stimulation (applied to each cortex and alternately to both cortices) on the locomotor output during the entire week following recovery of unsupported treadmill locomotion. Spontaneous treadmill locomotion was also tested biweekly throughout recovery (week 1-5), as well as functional responses to cortical stimulation and stimulation thresholds (Fig. 7). Kinematic analyses were automatically performed using DeepLabCut (Mathis, Mamidanna et al. 2018, Lecomte, Audet et al. 2021) and manually curated to correct misdetections. Obstacle avoidance performance was tested weekly, and analyses were double blinded. At the end of the experiments, after intracardiac perfusion, the spinal cord was extracted and processed for histological assessment of the spinal contusion. All procedures followed the guidelines of the Canadian Council on Animal Care and were approved by the Comité de Déontologie de l’Expérimentation sur les Animaux (CDEA, animal ethics committee) at Université de Montréal. All animals were included in the study.

### 2.2. Surgical procedures

All surgical procedures for electrode implantation or spinal lesions were conducted under general anesthesia and aseptic conditions. Animals were premedicated with Atravet (0.05 mg/kg), glycopyrrolate (0.01 mg/kg), and ketamine (10 mg/kg) by intramuscular administration. An endotracheal tube was then inserted to provide gaseous anesthesia (2% isoflurane in a mixture of 95% O2 and 5% CO2). In the first surgery, cats were implanted with intramuscular electrodes to record EMG activity from flexor and extensor hindlimb muscles on both sides, and they were also implanted with intracortical arrays to stimulate the hindlimb motor cortices during locomotion. The implanted muscles were semitendinosus (St; knee flexor and hip extensor), sartorius (Srt; hip flexor and knee extensor), vastus lateralis (VL; knee extensor), gastrocnemius lateralis and medialis (GL and GM; ankle extensors and knee flexors), as well as tibialis anterior (TA; ankle flexor). Electrodes were led subcutaneously to two 15-pinhead connectors secured to the cranium using acrylic cement. During the same surgery, a craniectomy was performed to expose the cruciate sulcus of both hemispheres and the dura was resected. The electrode array was stereotaxically inserted into the posterior bank of the cruciate sulcus that contains the hindlimb representation of the motor cortex (Nieoullon and Rispal-Padel 1976, Ghosh 1997, Bretzner and Drew 2005). One electrode array was inserted into layer V of each motor cortex (cortical area 4γ). In the first cat, a 5-channel array (P1 Technologies, USA) was implanted into the right motor cortex and 10 polyimide-coated stainless steel microwires (diameter: 50 µm, FineWire, USA) were implanted into the left motor cortex. The P1 array consisted of five individual electrode shanks (stainless steel, diameter: 250 µm) and a common ground. The other two cats were implanted with 32-channel arrays consisting of 4 shanks (silicon, length: 5 mm), spaced by 400 µm, each featuring 8 iridium active sites, spaced by 200 µm (NeuroNexus, USA). The most anteromedial site was lowered at 2 mm caudal to the cruciate sulcus and 2.5 mm lateral to the medial line. The cortex was covered with a hemostatic material (Gelfoam) and the arrays and EMG connectors were attached to the cranium with 8-10 screws and dental acrylic. Four to eight weeks after EMG and intracortical array implantation, a spinal contusion at T10 was achieved using a modified version of the Infinite Horizon Impactor model 0400 (Precision Systems and Instrumentations, USA) (Delivet-Mongrain, Dea et al. 2020). The impactor was mounted on the side of a spinal contention unit allowing the fixation of the T10 vertebra with clips to minimize movement during the application of the impactor tip. A force of 700 kdyne (7N), with a 15ms rise time, was maintained for 30 s through a 5 mm diameter flat circular tip applied on the dura. Heart rate and respiration were monitored throughout the surgeries. Twenty-four hours before each surgery, an antibiotic (Convenia, 8 mg/kg) was administrated subcutaneously. Before the end of the surgery, the analgesic buprenorphine (0.01 mg/kg) was administered subcutaneously. Additionally, a fentanyl patch (25 µg/h) was sutured to the skin to alleviate pain for ∼5 days. Gabapentin (10 mg bid) was also given for 3 days to alleviate pain if needed, after the implantation but not the spinal contusion surgery.

### 2.3. Behavioral assessments and analyses

Stepping patterns and skilled locomotion were assessed using the following two tasks: (1) treadmill without obstacles, and (2) treadmill with obstacles. Before the first surgery, all cats were trained to walk on a treadmill with positive reinforcement. They were not trained to avoid obstacles, but we occasionally introduced obstacles during the habituation period. During episodes of locomotion at the speed of 0.4 m/s, cats were recorded from the left and right sides with a digital video camera (120 Hz, Teledyne FLIR, USA). The kinematic analyses and the obstacle avoidance scoring were performed offline.

### Kinematic analysis

Reflective markers were placed over the iliac crest, the greater trochanter, the lateral malleolus, the metatarsophalangeal (MTP) joint, and at the tip of the fourth toe of the left and right hindlimbs. EMG signals were digitalized at 6 kHz (anti-aliasing filter at 45%) and filtered online (bandpass, 70-700 Hz) using a real-time BioAmp processor (Tucker–Davis Technologies, USA). Following data acquisition, we analyzed locomotor episodes of 10±2 consecutive step cycles using DeepLabCut (Mathis, Mamidanna et al. 2018, Lecomte, Audet et al. 2021) and manually curated to correct misdetections. A step cycle represents the time between two successive contacts of the same foot on the treadmill, beginning and ending with a foot strike. Hindlimbs are referred to as contralateral or ipsilateral to the stimulated cortex. The contralateral hindlimb is located on the opposite side of the stimulated cortex and the ipsilateral hindlimb is on the same side.

The following different kinematic parameters (indicated in italics) were used to evaluate the stepping abilities: The *step height* represented the maximum height (cm) of the foot during the swing phase. The *flexion velocity* corresponded to the maximum vertical speed (cm/s) reached by the foot during the swing phase. The *toe trajectory* represented the trajectory of the hindlimb’s toe (tip of the 4^th^ toe) during the step cycle. The *foot drag* was quantified as the time (expressed as a percentage of the swing phase) where the dorsal part of the distal phalanx of a given hindpaw dragged over the treadmill belt. The *lift* was the moment at which the foot took off from the treadmill belt and the *contact* was the moment at which it contacted the belt. In order to normalize a step cycle, we used the contact of the hindlimb ipsilateral to the stimulated cortex as a reference. The *lift phase* corresponded to the moment at which the lift of the contralateral hindlimb occurred in a normalized step cycle. The *lift phase variability* was the standard deviation of the lift phase inside a locomotor episode (10±2 step cycles). The *contact phase* corresponded to the moment at which the contact of the contralateral hindlimb occurred in a normalized step cycle. The *stimulation onset phase* corresponded to the moment at which the stimulation burst began during a normalized step cycle. The *stimulation precision* indicated the % at which the stimulation onset phase began in a 20% phase interval of the step cycle (expected stimulation time window) where the highest step height increase (in the intact state) and dragging reduction (after SCI) were achieved. This interval included the lift of the contralateral hindlimb. The *stance length* was the distance (cm) traveled by the hindlimb during the stance. The *foot position at contact* was the distance (cm) between the toe and the hip (vertical projection of the hip’s position) at the hindlimb’s contact. This quantity indicated the extent of forward movements during a locomotor episode. The s*tep cycle duration* represents the time (s) between two consecutive contacts of the same foot on the treadmill, whereas *swing duration* referred to the time (s) between toe-off and foot contact. The *stance duration* referred to the time between foot contact and toe-off. The *trajectory modulation* comprised the time series representing the average effect of a given stimulation condition on the swing trajectory. They are defined as the difference between spontaneous trajectories and modulated trajectories, in polar coordinates. The center of the polar coordinate system was set at the center of the segment connecting the points where the foot takes off the ground and strikes the ground. The *centers of modulation* of the foot trajectories are vectors, which indicate the centroid (in polar coordinates) of trajectory modulations, as depicted in Fig. 4E. Multivariate analysis consisted of a PCA dimensionality reduction of each trajectory modulation series into the 2D space of the two principal components. Each projected point represents the average modulation obtained with a given stimulation condition.

### Obstacle avoidance analysis

Following a paradigm established by Drew (1993), an obstacle (height: 5 cm; anteroposterior width: 5 cm; mediolateral width: 35 cm) was placed on the treadmill belt. Five-minute videos were captured weekly and analyzed in double blind to evaluate the ability to step over the obstacle without touching it in the intact state and after spinal contusion.

### 2.4. Phase-coherent ICMS during locomotion

Biphasic (cathodic first) 330 Hz stimulations were delivered with a Tucker-Davis Technologies (USA) stimulator during treadmill locomotion. A simple EMG pattern recognition algorithm was used to detect the gait phases (Bonizzato and Martinez 2021). Each time the filtered and rectified signal from a selected muscle (gastrocnemius medialis ipsilateral to the stimulated cortex) reached a manually selected threshold, a gait synchronization event was detected, indicating the onset of muscle activity. For each detection, a refractory period of 700ms was imposed to prevent eliciting triggers within a single EMG burst. Synchronization triggered the delivery of a phasic stimulation to a selected electrode in the array (100ms burst, 330 Hz frequency, biphasic, cathodic first, 200 µs/phase, and 50 µs phase interval). For intact cats, 4-8 electrode sites per cortex were chosen for recordings after visually determining that they elicited different types of movements. After contusion, 3-6 electrode sites per cortex were chosen for their ability to produce a flexion movement.

#### 2.4.1. Phase-coherent uni-cortical stimulation

In intact cats and SCI cats, we tested the effects of varying stimulation parameters on the evoked locomotor output. During locomotor episodes, uni-cortical stimulation was delivered with one parameter being modulated while the others remained constant. Uni-cortical stimulation is defined as stimulation that was delivered to only one motor cortex during a locomotor episode. The modulated parameters were the timing, the burst duration, the amplitude, and the site of stimulation. When the parameters were kept constant, we used: a sub-maximal amplitude (high amplitude), a 200ms delay after synchronization event detection (corresponding to the contralateral hindlimb’s lift), and a 100ms burst duration. For the study of timing, burst duration, and amplitude, diverse stimulation channels within each cortical array were selected among those with the lowest current threshold for evoking movement.

#### Timing

Different delays between the detection of a synchronization event (onset of left or right gastrocnemius medialis activity) and the delivery of the stimulation (0, 100, 200, 300, and 400ms from each of the two synchronization events) were assessed. Each delay corresponded to a different gait cycle timing and covered the entire step cycle of the animal. Each delay was tested randomly. Every delay (in ms) between the synchronization event and the onset of stimulation was the same, but the exact normalized phase of the step cycle at which the onset of stimulation occurred varied. The data was linearly interpolated using Interp1 (MATLAB 2019b) and was averaged across all tested sites and all cats.

#### Burst duration

Both before and after SCI, the effect of burst duration on the evoked motor response was tested using four different stimulation train durations: 50, 100, 150, and 200ms.

#### Amplitude

The stimulation amplitude was modulated within a “functional range” of amplitudes. During ongoing locomotion, we stimulated with increasing amplitudes to find the threshold (i.e. the smallest amplitude evoking a visible response; intact: 10-75 µA, post-SCI: 50-300 µA), which corresponded to the beginning of the range. We then continued to increase the amplitude until the maximum comfortable value was reached (intact: 30-250 µA, post-SCI: 100-500 µA). The range was divided into five amplitudes (threshold, low, medium, high, and maximum) for intact cats and three amplitudes (threshold, medium, and maximum) for SCI cats. The medium value indicates the average between the threshold value and the maximum value (50% of the range). The low and high values represent 25% and 75% of the functional range, respectively. The different amplitudes were randomly tested for every stimulation site.

#### 2.4.2. Phase-coherent bi-cortical stimulation

We also tested whether stimulation applied alternately to the left and right motor cortex during walking could bilaterally modulate hindlimb locomotion.

The stimulation was delivered to both cortical hemispheres alternately and phase-coherently during treadmill locomotion (delay: 200ms after synchronization event detection, burst: 100ms). The stimulus amplitude applied to each cortex was independently selected and we tested all combinations (a full 2D matrix of stimulation amplitude options, left vs right cortex: 4×4 for intact cats and 3×3 for SCI cats). In intact cats, four different amplitudes were used: spontaneous (stimulation off), threshold, medium, and maximum. In SCI cats, only three amplitudes were tested: spontaneous, medium, and maximum.

### 2.5. Evaluation of the lesion size

At the end of the experiments, the animals were deeply anesthetized with ketamine (Ketaset; 10 mg/kg; intramuscular) and administered a lethal dose of pentobarbital (Euthanyl; 120 mg/kg; intravenous). The animals were perfused transcardially with a solution of 0.2% heparin in 0.1 M phosphate-buffered saline (PBS; pH 7.4), followed by 4% paraformaldehyde (PFA) in 0.1 M PBS (pH 7.4). The spinal cord was extracted and postfixed for 24 h in a solution of 16% PFA in 0.1 M PBS. The tissue was then cryoprotected in a solution of 30% sucrose in 0.1 M PBS. The spinal cord was frozen, and 40 µm thick coronal sections centered on the spinal lesion were taken for histological examination. Every third section was mounted on slides and stained with Luxol fast blue (0.1%, Sigma, USA S3382) to visualize myelin in the spinal white matter and cresyl violet (0.5%, Alfa Aesar, USA J64318) to visualize cell bodies in the spinal gray matter. Bright-field microscopy images were taken at 4X (Olympus BX63) and analyzed using cellSens software (Olympus, Japan). Lesion extent was quantified by evaluation of the percentage of damaged and intact tissue of the cord observed in the coronal plane. Each image area was assigned to one of three categories [cavity, damage, intact], and the number of pixels belonging to each category was counted.

### 2.6. Quantification and statistical analyses

All data are presented as the mean values ± SEM. All statistical analyses were performed using GraphPad Prism 6 or MATLAB software. The normal distribution of the data was assessed using the Anderson-Darling test. We first performed a one-way repeated measures ANOVA. The Fisher LSD test was used as a post-hoc for multiple comparisons. When multiple comparisons were conducted, the Bonferroni method was used to re-establish the level of significance and control for false positives. The tests were one-sided, because our hypotheses were strictly defined toward the direction of motor improvement. The statistical significance threshold was set at P=0.05.

## 3. Results

We designed a neuroprosthetic platform whereby ongoing locomotor phases were monitored through real-time online processing of electromyographic (EMG) activity from hindlimb muscles (Fig. 1). Hindlimb contacts with the ground during gait (“foot-strikes”) were detected by pattern recognition of gastrocnemius muscle activity. ICMS was triggered at a fixed delay following foot-strikes, corresponding to the expected lift of the hindlimb contralateral to the stimulated cortex. Before and after a spinal cord contusion, we first characterized the effects of multiple stimulation parameters on kinematics, including timing, stimulus duration, amplitude, and site of stimulation. We explored ICMS controllability over a repertoire of varied hindlimb movements. Before and after a spinal cord contusion we also tested whether bi-cortical ICMS could be used to control bilateral gait performance by enforcing alternated patterns of stimulation, coherent with each hindlimb’s movement. After SCI, we determined what stimulation parameters maximally alleviated foot drop. We found that the re-emergence of cortical neuroprosthetic control of movement and spontaneous weight-supporting locomotion occurred synchronously in all cats.

**Fig. 1.**
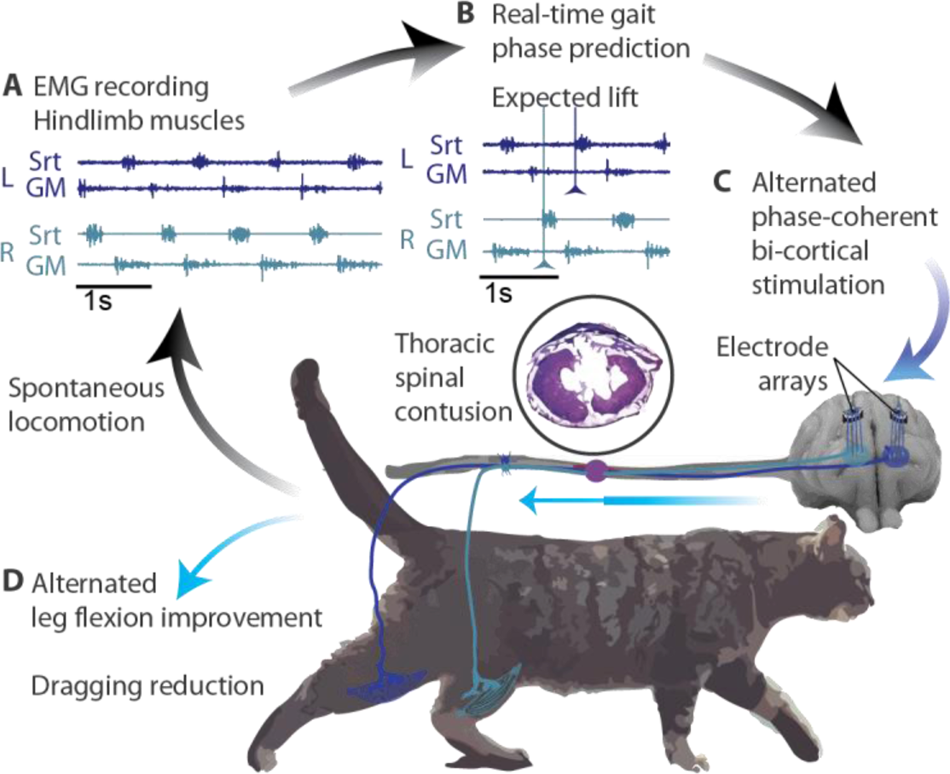
Bi-cortical neuroprosthesis design. **(A)** During treadmill locomotion, electromyographic activity was recorded from flexor and extensor muscles from both hindlimbs. **(B)** EMG activity was analyzed in real-time to predict the occurrence of each hindlimb’s foot lift. **(C)** Lift prediction triggered ICMS delivery to hindlimb motor cortices. **(D)** ICMS applied to the right motor cortex enhanced leg flexion of the left hindlimb and conversely, modulating bilateral locomotor output. L: left, R: right, Srt: sartorius, GM: gastrocnemius medialis, and s: seconds.

### 3.1. Phase-coherent ICMS modulated contralateral hindlimb kinematics in healthy cats

The modulation of hindlimb motor output enabled by ICMS was investigated in n=3 intact cats. During treadmill walking, ICMS was delivered through an electrode selected from an array implanted in the left or right hindlimb motor cortex (Fig. 2A). Using EMG pattern recognition, timely delivery of the stimulation was triggered within the expected time window, which encompassed the lift of the contralateral hindlimb, with 83.4% precision (Fig. S1A). Changes in hindlimb trajectory and locomotion were assessed while varying either the timing, duration, or amplitude of the stimulation.

**Fig. 2.**
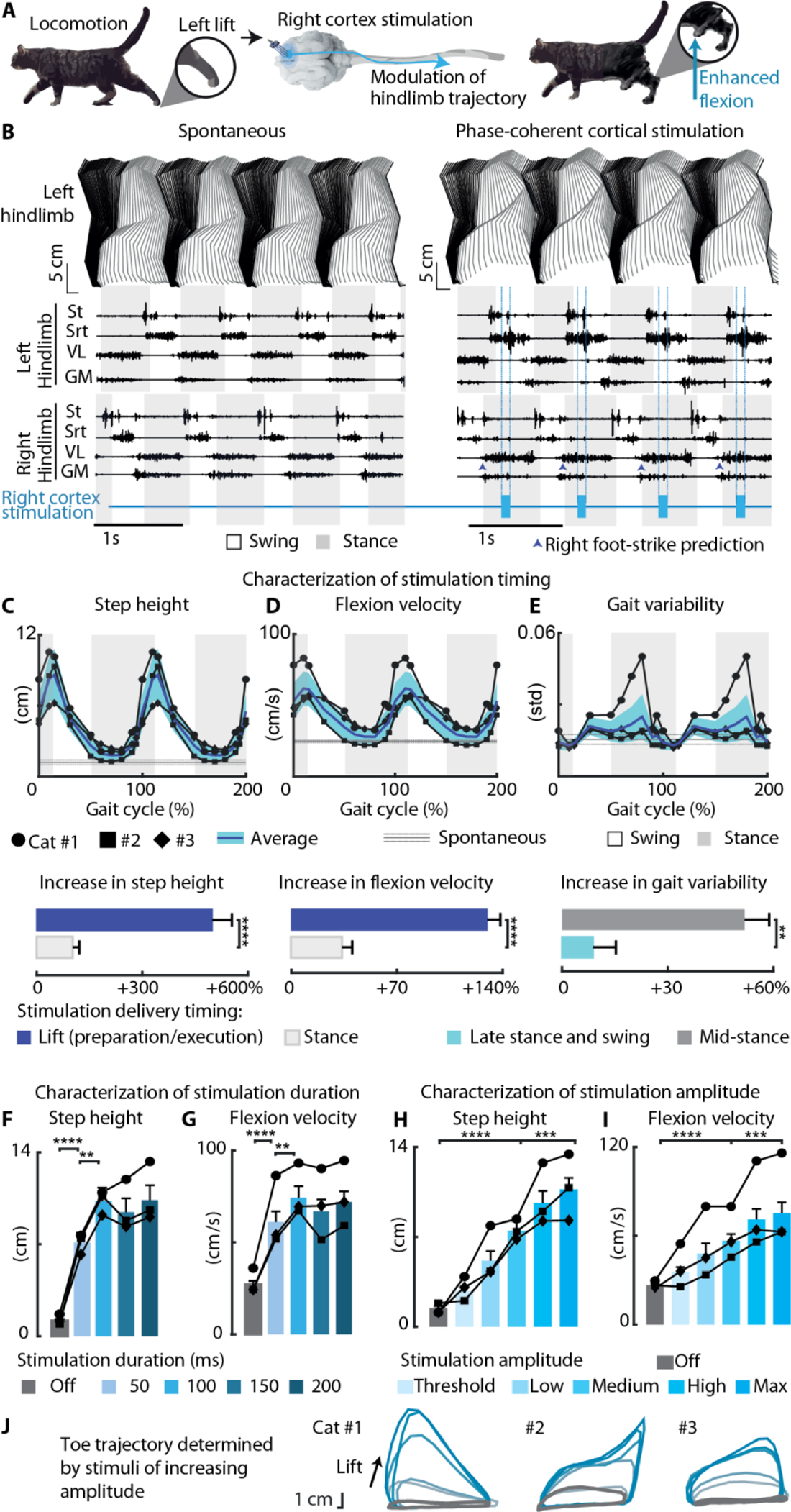
Uni-cortical stimulation modulated contralateral hindlimb kinematics in intact cats. **(A)** Schematic representation of uni-cortical neurostimulation: right cortex stimulation modulated left hindlimb flexion through descending projections. **(B)** Stick diagram and EMG activity during spontaneous locomotion and phase-coherent ICMS delivered to the right cortex. Changes in **(C)** step height, **(D)** flexion velocity, and **(E)** gait variability (n=17 sites, 3 cats) as a function of stimulus delivery timing along the gait cycle. Changes in **(F)** step height and **(G)** flexion velocity (n=16 sites, 3 cats) as a function of burst duration. Increasing ICMS amplitude linearly modulated **(H)** step height, **(I)** flexion velocity (n=17 sites, 3 cats), and **(J)** toe trajectories. p: *<0.05; **<0.01; ***<0.001; ****<0.0001. St: semitendinosus, Srt: sartorius, VL: vastus lateralis, and GM: gastrocnemius medialis.

Stimulus timing played a critical role in neuromodulation efficacy. When delivered at the beginning of the contralateral hindlimb swing, ICMS produced an enhanced contralateral hindlimb flexion in comparison to spontaneous walking (Fig. 2B). The largest changes in step height (p<0.0001, up to an average of +575±78% of spontaneous walking) and flexion velocity (p<0.0001, up to +141±20% of spontaneous walking) were obtained when ICMS was delivered between the contralateral hindlimb lift preparation (late stance) and execution (early swing) (Fig. 2C-D). Conversely, ICMS delivered during the contralateral hindlimb’s stance phase did not modulate step height and flexion velocity in comparison to spontaneous locomotion (p>0.05). ICMS delivered in the middle of the contralateral hindlimb’s stance phase disrupted locomotion (Fig. 2E) and increased the lift phase variability (p<0.05, up to +62±26%). ICMS had no effect on the ipsilateral hindlimb’s kinematics, even when delivered during its swing phase (Fig. S2A-B). We defined ICMS to be “phase-coherent” with locomotion when delivered in phase with the contralateral hindlimb lift preparation.

We then evaluated the impact of phase-coherent ICMS duration on hindlimb flexion. The largest increase in step height (p<0.0001, +915±152% of spontaneous walking) and flexion velocity (p<0.0001, +178±21% of spontaneous walking) were obtained with a 100ms stimulation duration (Fig. 2F-G). Kinematic modulation plateaued for longer durations. A 100ms duration was considered optimal and was consequently used in all experiments.

We finally examined the effects of varying phase-coherent ICMS amplitude on hindlimb flexion, revealing a high-fidelity proportional control of motor output. Stimulation amplitude featured precise control over contralateral toe trajectories (Fig. 2H-J) but had no impact on ipsilateral hindlimb flexion (Fig. S2C-D). Contralateral step height (p<0.0001, up to +720±91% of spontaneous walking, fit r^2^: 90±8%) and flexion velocity (p<0.0001, up to +179±21% of spontaneous walking, fit r^2^: 85±10%) modulated linearly with increasing stimulation amplitudes (Fig. 2H-I).

### 3.2. Phase-coherent ICMS modulated contralateral hindlimb kinematics after SCI

Thoracic spinal cord contusion produced complete acute bilateral hindlimb paralysis. Two to three weeks after SCI, all n=3 cats exhibited weight-supported bilateral hindlimb locomotion, despite severe dragging (Fig. 3A-B). We tested the immediate effects of phase-coherent ICMS during treadmill walking. Timely delivery of the stimulation to selected electrodes in the right or left motor cortex was achieved with 88.2% precision (Fig. S1C).

**Fig. 3.**
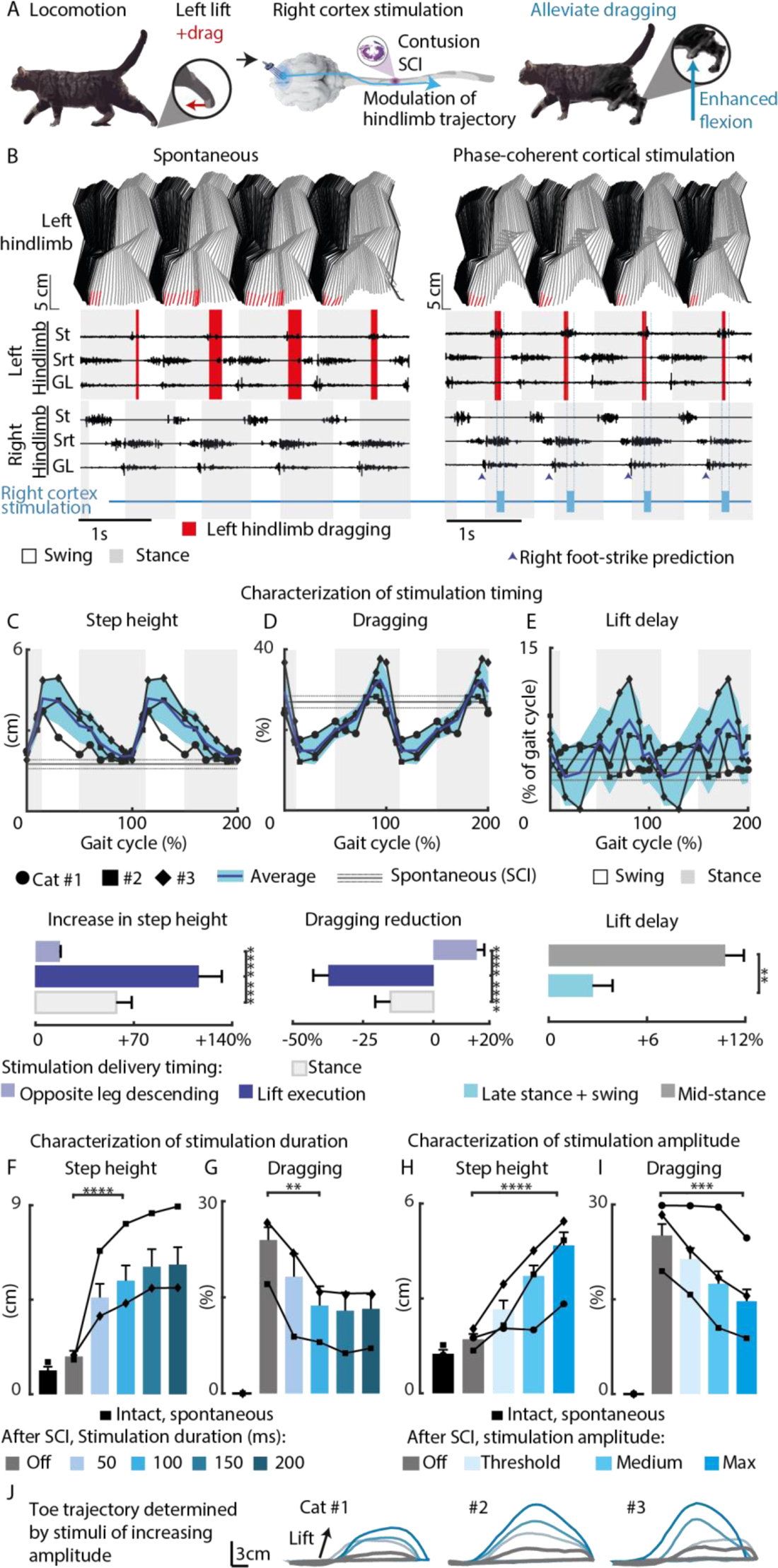
Uni-cortical stimulation alleviated contralateral hindlimb deficits after SCI. **(A)** Schematic representation of uni-cortical neurostimulation following thoracic spinal cord contusion. ICMS modulated contralateral hindlimb locomotion through spared descending nerve fibers. **(B)** Stick diagram and EMG activity during spontaneous locomotion and with phase-coherent ICMS delivered to the right cortex. When applied during swing execution, ICMS enhanced **(C)** step height and reduced **(D)** dragging. When applied during mid-stance, ICMS modified the **(E)** lift delay of the contralateral hindlimb (n=15 sites, 3 cats). The burst duration modulated the **(F)** step height, and reduced **(G)** dragging of the contralateral hindlimb (n=7 sites, 2 cats). The stimulation amplitude linearly increased the **(H)** step height, decreased **(I)** dragging (n=15 sites, 3 cats), and modulated the **j** toe trajectories. p: *<0.05; **<0.01; ***<0.001; ****<0.0001. St: semitendinosus, Srt: sartorius, and GL: gastrocnemius lateralis.

Stimulus timing played a critical role in neuromodulation efficacy after SCI. Phase-coherent ICMS successfully enhanced contralateral hindlimb flexion, which, in turn, alleviated dragging in all tested animals (Fig. 3B). An increase in step height (p<0.0001, up to +140±16% of spontaneous walking; Fig. 3C) and reduction of dragging (p<0.001, up to −45±5% of spontaneous walking; Fig. 3D) were maximal when stimulation was delivered during contralateral swing execution. Conversely, stimulation delivered during contralateral swing preparation did not significantly increase step height (p>0.05, +19±7% of spontaneous walking post-SCI, Fig. 3C) and was associated with a non-significant increase in dragging (p>0.05, up to +20±9% of spontaneous walking, Fig. 3D). When stimulation was delivered in the middle of the contralateral stance phase, it disrupted the step cycle by delaying the beginning of the lift phase (p<0.05, up to +13±5% of spontaneous walking, Fig. 3E). As observed in intact cats, ICMS did not affect ipsilateral hindlimb trajectory, independently of the stimulation timing (Fig. S2E-F). Hence, in SCI cats, we defined stimulation as “phase-coherent” with locomotion when delivered during the execution phase of the contralateral hindlimb lift.

We then evaluated the effects of phase-coherent ICMS duration on hindlimb flexion. Consistently with the results obtained in the intact state, a 100ms stimulation train produced a significant increase in step height (p<0.0001, 241±62% of spontaneous walking, Fig. 3F), which, in turn, decreased dragging (p<0.01, −42±10% of spontaneous walking, Fig. 3G). Since further increasing the stimulus duration only generated minimal gains in movement modulation, stimulus duration was maintained at 100ms in all subsequent experiments.

We finally examined the effects of varying phase-coherent ICMS amplitude on hindlimb flexion after SCI (Fig. 3H-J). As observed in the intact state, ICMS amplitude modulated contralateral leg trajectory (Fig. 3J), with no impact on ipsilateral hindlimb flexion (Fig. S2G-H). Contralateral step height (p<0.0001, up to +202±31% of spontaneous walking, fit r^2^: 78±30%, Fig. 3H), and dragging reduction (p<0.0001, up to −42±6% of spontaneous walking, fit r^2^: 68±25%, Fig. 3I) modulated linearly with increasing stimulation amplitudes.

### 3.3. Phase-coherent ICMS produced diverse movement synergies in intact cats that were lost after SCI

ICMS produced a variety of motor outputs in intact cats. When selecting different electrode sites within the hindlimb motor cortex representation, we were able to impose qualitative changes to hindlimb trajectories during leg swing. Six distinct motor programs were recruited across intact animals, and the associated gait trajectories were characterized under increasing stimulation amplitudes (Fig. 4A-B, Fig. S3, Movie S1). Each movement remained qualitatively similar with varying stimulation amplitudes within the same electrode, while varying in the extent of trajectory modulation. Switching between different stimulation electrodes, we obtained trajectory modulations consisting of backward flexions, abductions, forward flexions, qualitatively different upward flexions, including ones appearing ‘natural’ (i.e., a smooth, curved progression of the limb up and forward), as well as some associated with a co-contraction of flexor and extensor muscles. In cat #3, we also identified a swing-stop control (Fig. S4A), whereby stimulation delivery shortened the swing movement, resulting in curtailment of the step (Fig. S4B-C). We repeated the same characterization after SCI and found that this evoked movement diversity was absent. Across all cats, stimulation resulted in upward-directed modulation of hindlimb flexion, for all sites and stimulation amplitudes (Fig. 4C-D).

We performed an analysis of movement trajectories by (1) comparing the main direction of swing modulation (‘centers of modulation’) (Fig. 4E) and (2) performing multivariate analysis (principal component analysis, PCA) of gait trajectories (Fig. 4F) across all motor programs. PCA grouped the identified motor programs in separate zones of the principal component space, with movements evoked in intact cats scattered throughout the PC space. In contrast, all movements evoked after SCI were concentrated in only one area.

**Fig. 4.**
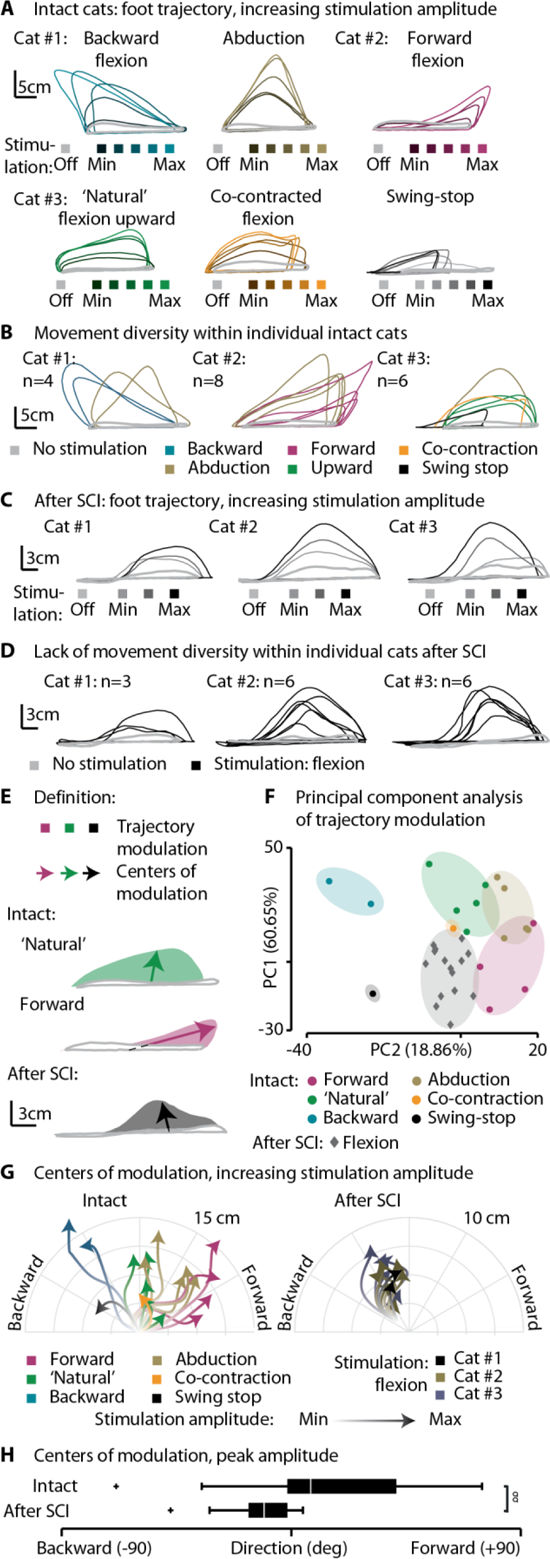
Uni-cortical stimulation triggered a variety of motor synergies in intact cats and a single stereotyped lift movement after SCI. (A) Six different movements evoked by varying stimulation electrodes across three intact cats. For each stimulation electrode, superimposed trajectories were color-coded for the tested range of stimulation amplitudes. (B) The peak modulations of each tested electrode were superimposed for each cat. (C) Movements evoked by cortical stimulation across three SCI cats. For each stimulation electrode, superimposed trajectories were color-coded for the tested range of stimulation amplitudes. (D) The peak modulations of each tested electrode were superimposed for each cat. (E) Schematic description of the trajectory modulation assessments (difference in foot trajectory with and without stimulation) and centers of modulation (centroid of trajectory modulations). (F) Principal component analysis of trajectory modulations. The data were color-coded by visual qualification of evoked movements. (G) Direction of modulation centers for intact and SCI cats. The arrows follow increasing stimulation amplitudes. (H) Distribution of modulation direction centers under peak stimulation amplitudes. ^σσ^: p<0.01, the symbol σ indicates a statistical test of the distributions’ variances. PC: principal component.

Analysis of the trajectory in intact cats displayed a large repertoire of directional swing movement modulations. After SCI, all modulations were pointed within a restrained portion of the polar diagram (Fig. 4G-H, p<0.01: two-sample two-sided F-test for equal variances).

### 3.4. Phase-coherent bi-cortical stimulation bilaterally modulated hindlimb trajectories in intact and SCI cats

Since phase-coherent ICMS delivered to one cortex manipulated contralateral hindlimb movements, we hypothesized that alternately stimulating both motor cortices would allow bilateral control of gait trajectories. Using EMG pattern recognition, timely delivery of the stimulation was triggered with 97.6% and 98.1% precision in intact and SCI cats, respectively (Fig. S1B-D). Before and after contusive SCI, phase-coherent ICMS (100ms) was delivered alternately to the left and right motor cortex during treadmill walking in n=3 cats (Fig. 5A and 6A). Delivery of bi-cortical stimulation induced an alternated increase of hindlimb flexion in intact cats (Fig. 5B, Movie S2). After SCI, phase-coherent bi-cortical stimulation immediately improved the bilateral locomotor pattern, enhancing hindlimb flexion, which, in turn, alleviated dragging deficits (Fig. 6B, Movie S3). The beneficial effects of ICMS disappeared when the stimulation was discontinued.

**Fig. 5.**
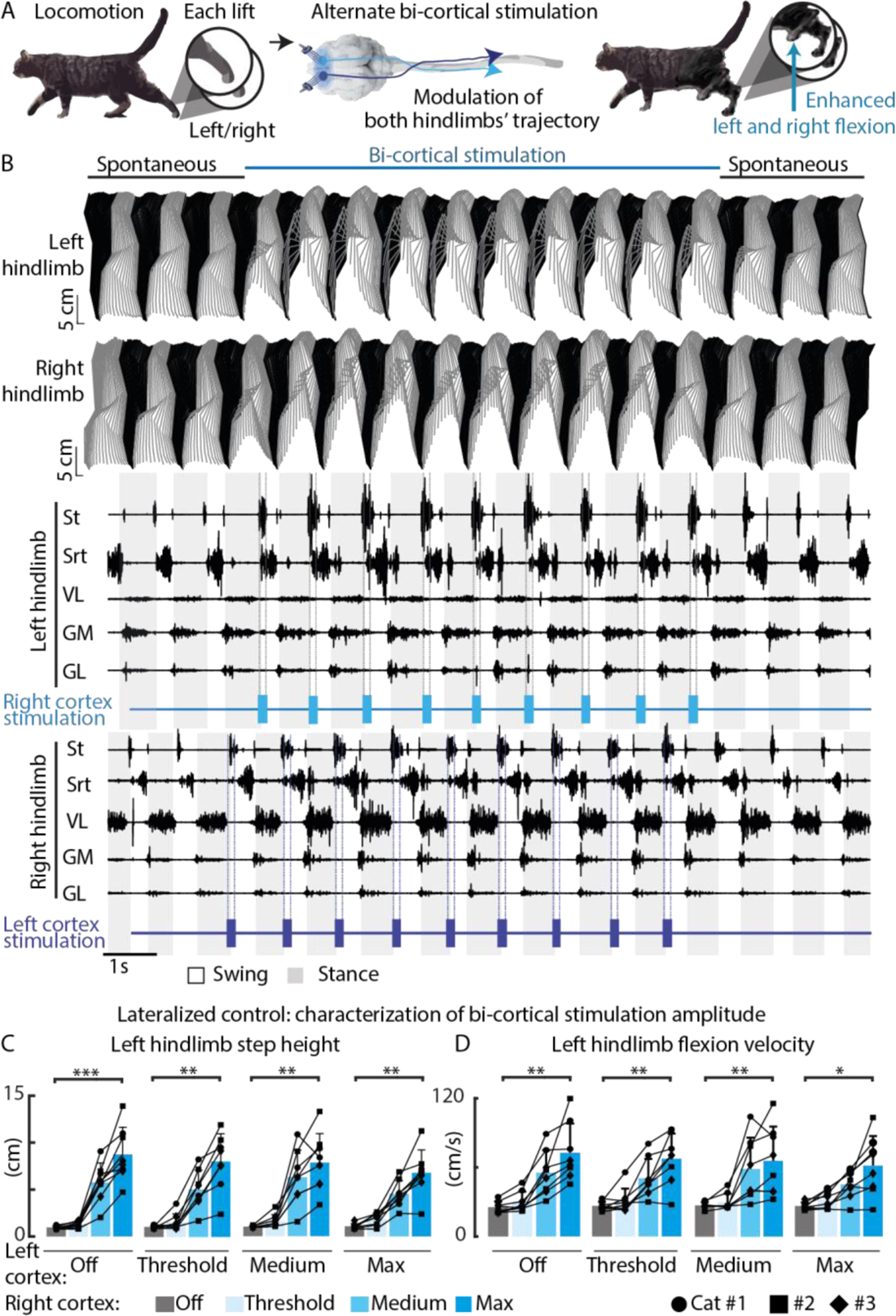
Phase-coherent bi-cortical stimulation modulated bilateral gait kinematics in intact cats. **(A)** Schematic representation of bi-cortical neurostimulation. Both cortices were stimulated alternately and in phase coherence with locomotion. **(B)** Stick diagram and EMG activity during spontaneous walking with or without bi-cortical stimulation. During bi-cortical stimulation, the **(C)** left hindlimb step height and **(D)** left hindlimb flexion velocity linearly modulated with increasing ICMS amplitude applied to the right cortex, independently of the stimulation amplitude delivered to the homologous cortex. p: *<0.05; **<0.01; ***<0.001. St: semitendinosus, Srt: sartorius, VL: vastus lateralis, GM: gastrocnemius medialis, and GL: gastrocnemius lateralis.

**Fig. 6.**
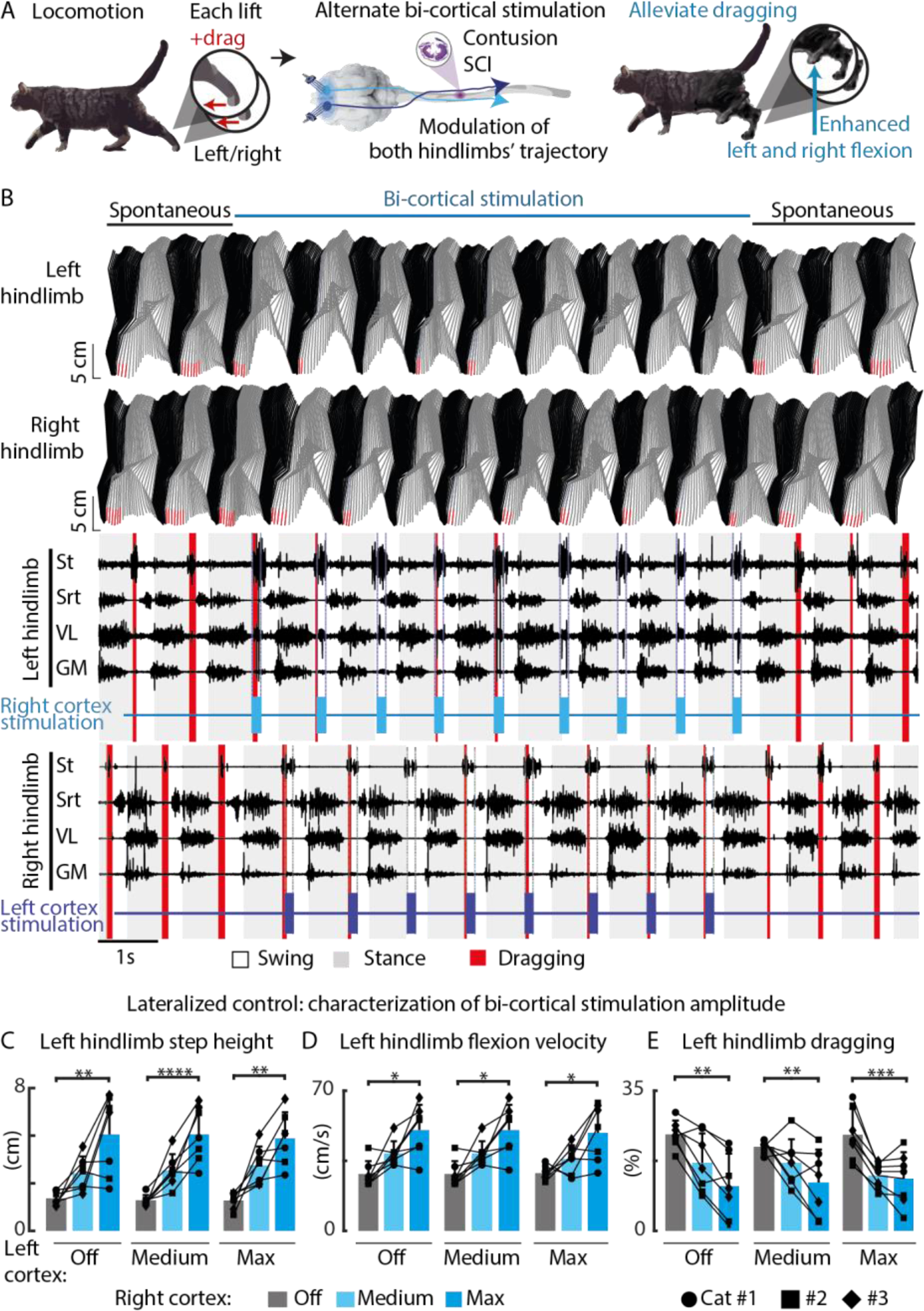
Phase-coherent bi-cortical stimulation bilaterally modulated gait kinematics and reduced foot drop deficits after SCI. **(A)** Schematic representation of bi-cortical neurostimulation after an incomplete contusive SCI. Both cortices were stimulated alternately and in phase coherence with locomotion. **(B)** Stick diagram and EMG activity during spontaneous walking, with or without bi-cortical stimulation. During bi-cortical stimulation, the **(C)** left hindlimb step height, **(D)** left hindlimb flexion velocity and **(E)** left hindlimb dragging linearly modulated with increasing ICMS amplitudes applied to the right cortex, independently of the stimulation amplitude delivered to the homologous cortex. p: *<0.05; **<0.01; ***<0.001. St: semitendinosus, Srt: sartorius, VL: vastus lateralis, and GM: gastrocnemius medialis.

Next, we investigated the extent to which the specific cortical control of contralateral hindlimb movements is independent from interactions with the activated homologous cortex. Thus, we combined the stimulation delivered to the right and left motor cortex, independently selecting the stimulus amplitude on each cortex and testing all combinations (a 2D matrix of stimulation amplitude options: 4×4 for intact cats and 3×3 for SCI cats). In intact cats, the contralateral step height (p<0.01, up to +681±104% of spontaneous walking, fit r^2^: 90±1%, Fig. 5C) and flexion velocity (p<0.01, up to +159±31% of spontaneous walking, fit r^2^: 85±2%, Fig. 5D) modulated linearly with increasing amplitudes. Stimulation of the homologous cortex did not interfere with the specific cortical stimulation effects on contralateral hindlimb movements (Fig. S5A-B).

After SCI, increasing stimulation amplitudes produced a linear increase in contralateral hindlimb step height (p<0.01, up to +234±52% of spontaneous walking post-SCI, fit r^2^: 84±5%, Fig. 6C) and flexion velocity (p<0.05, up to +82±19% of spontaneous walking post-SCI, fit r^2^: 75±3%, Fig. 6D), which, in turn, produced a linear decrease in dragging (p<0.001, up to −48±7% of spontaneous walking post-SCI, fit r^2^: 76±3, Fig. 6E). The specific effect of cortical stimulation on contralateral hindlimb movements was independent from the level of stimulation of the homologous cortex (Fig. S5C-E).

### 3.5. The return of locomotion after SCI was time-locked with descending drive recovery

Contusive SCI are characterized by the interruption of variable pathways and pronounced secondary damage, including cavity formation (Delivet-Mongrain, Dea et al. 2020). In this study, the spinal lesions exhibited central cavitations and surrounding intact tissue that contained residual descending motor pathways, including pyramidal and non-pyramidal tracts (Fig. 7A, D, and G).

**Fig. 7.**
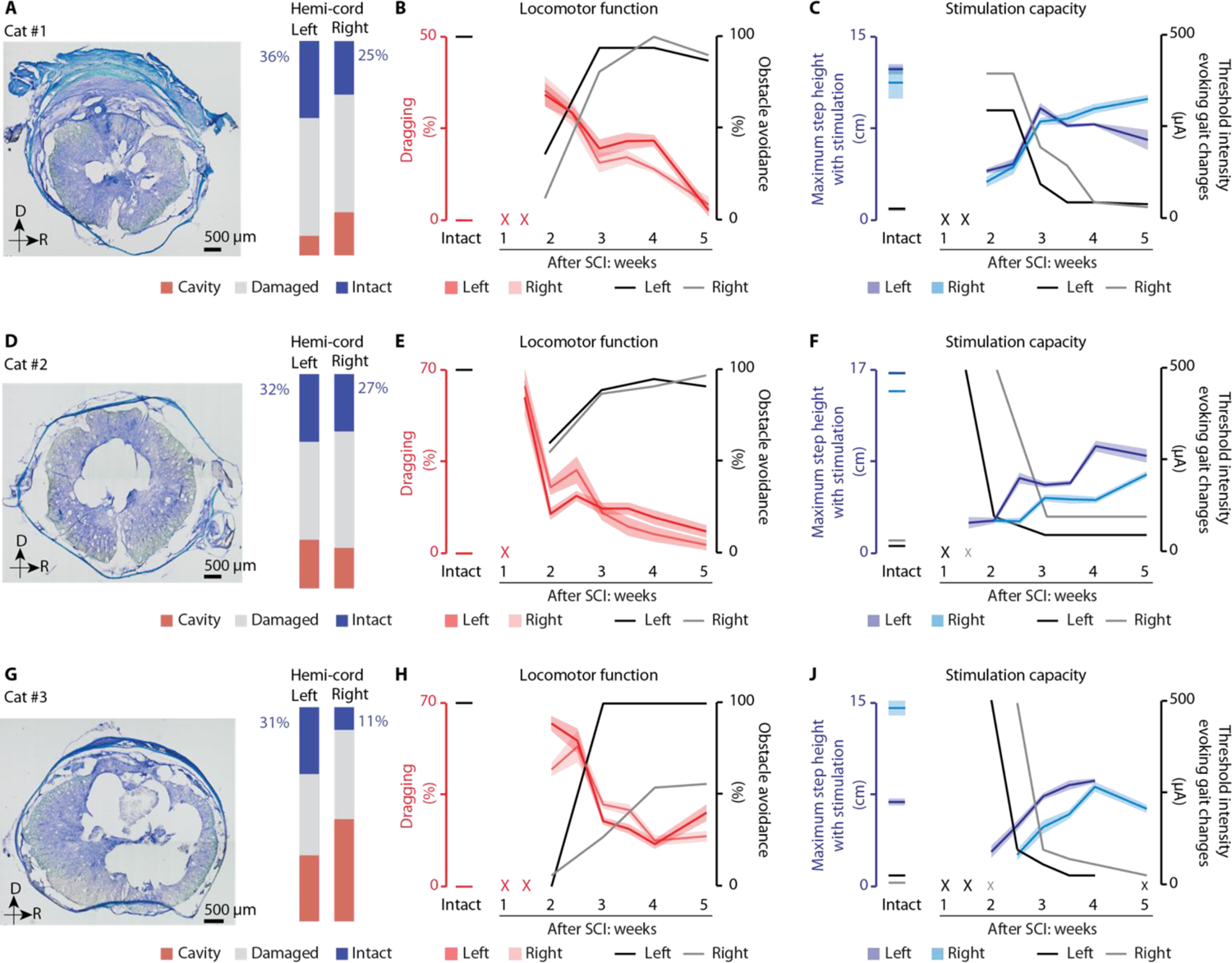
The return of locomotion was time-locked with the re-emergence of cortically-induced gait modulation. Data for cat #1: **(A)** Coronal sections of the lesion epicenter were stained with Luxol fast blue and cresyl violet. The contusion epicenter was imaged at 4X magnification to assess the extent of the lesion. Scale: 500 µm. D: dorsal. R: right. **(B)** After SCI, cats displayed bilateral hindlimb dragging during spontaneous walking. Dragging decreased over time, while voluntary motor control, as quantified by the obstacle avoidance task, increased over time. Red X: cat was unable to express weight-supported locomotion at the indicated timepoints. **(C)** Stimulation capacity assessed in terms of maximum step height obtained with cortical stimulation and minimum threshold stimulation amplitude, across all electrodes. The maximum step height increased over time, while the thresholds decreased. Black X: no electrode produced kinematic responses at this time point for both legs; the lack of effects is limited to one leg when the X value is smaller (shade indicates limb). **(D-F)** Data for cat #2 and **G-I** data for cat #3 as described in **A-C** above. In all line plots, the lone datapoint on the left represents the respective values for intact cats.

The three injury profiles were associated with distinct timelines of locomotor recovery. Cat #2 first recovered weight-supported treadmill locomotion 1.5 weeks after SCI and displayed moderate (>50%) obstacle avoidance performance by week 2 (Fig. 7E). Cat #1 was first able to walk unsupported at week 2, with low (<50%) obstacle avoidance performance (Fig. 7B). Cat #3 also recovered weight-supported locomotion at week 2 but exhibited severe dragging deficits (>50%) and inability to avoid obstacles (<5%), that persisted for half a week (Fig. 7H). Consistent with higher injury lateralization towards the right of the spinal cord, the left hindlimb performance on the obstacle avoidance task was almost two-fold improved over the right one. Over the course of 5 weeks after SCI, all n=3 cats partially recovered both hindlimb clearance during treadmill locomotion and obstacle avoidance capacity.

The three injury profiles were also associated with distinct timelines of stimulation output re-expression. Cat #2 first re-expressed cortically evoked hindlimb movements at 1.5 weeks (left hindlimb) and 2 weeks (right hindlimb) after SCI and displayed lower ICMS thresholds (<100 µA) by week 2 and 3, respectively (Fig. 7F). Cat #1 first expressed cortical modulation of gait at week 2, with high (>250 µA) ICMS thresholds (Fig. 7C). Cat #3 also started responding to cortical stimulation at week 2 on the left side, with high ICMS thresholds (>250 µA), which rapidly decreased in 3 days (Fig. 7I). Again, consistent with its SCI profile, the right hindlimb responses of cat #3 to stimulation and the threshold curves of the contralateral cortex were time-shifted by 3 days. All n=3 cats re-expressed increasingly stronger cortically-controlled gait modulation, with ICMS thresholds continuously decreasing over the course of 5 weeks after SCI.

## 4. Discussion

We developed a bi-cortical neuroprosthesis to alleviate the severe bilateral locomotor deficits produced by contusive SCI. To independently control the trajectory of each hindlimb after SCI, neurostimulation was applied alternately to the left and right motor cortex in phase coherence with locomotion. In healthy cats, cortical stimulation elicited a variety of motor programs. In contrast, after SCI, the motor programs switched to a single stereotyped vertical flexion movement, which counteracted foot drag. Even in cats exhibiting an asymmetrical gait pattern, an optimal setting of stimulation parameters allowed for the restoration of well-organized bilateral locomotion. Cortical neuroprosthetic controllability re-emerged in synchrony with spontaneous recovery of locomotion.

The motor cortex plays a major role in voluntary control of locomotion (Drew, Jiang et al. 2002). In cats and humans, damage to the motor cortex projections that are linked to the spinal circuits is associated with locomotor impairments (Jiang and Drew 1996, Barthelemy, Willerslev-Olsen et al. 2010). As previously observed in intact cats and rats, ICMS delivered during locomotion produces phase-dependent changes in locomotor activity and controls contralateral hindlimb trajectories (Bretzner and Drew 2005, Bonizzato and Martinez 2021, Fortier-Lebel, Nakajima et al. 2021, Massai, Bonizzato et al. 2021). In intact and hemisected rats, phase-coherent cortical stimulation immediately enhanced contralateral hindlimb locomotor kinematics (Bonizzato and Martinez 2021). Given its role in modulating locomotor output, the motor cortex is a target of choice for neuromodulation interventions intended to control limb movement during gait. The major goal of this study was to select and enhance descending motor commands involved in locomotor control of both hindlimbs by bilaterally targeting the motor cortex.

In clinical settings, most SCIs are caused by closed traumas with displaced vertebral fractures and associated spinal contusion (RHI 2018, N.S.C.I.S.C 2019), which in most cases affect both legs and may induce asymmetrical locomotor deficits. In our cat model of contusive SCI, alternate bi-cortical neurostimulation had independent and limb-specific effects, allowing for the control of bilateral gait patterns by tuning optimal stimulation amplitude settings, even in cats that displayed an asymmetrical locomotor pattern.

Stimulation timing was essential to obtain an enhanced and integrated movement as well as to reduce foot drag after SCI. The optimal timing was synchronous with the contralateral hindlimb lift, which we defined as “phase-coherent” stimulation. Phase-coherent stimulation is an emerging paradigm in neuromodulation of locomotion. It is superior to continuous neurostimulation in animal and human studies involving spinal cord stimulation (Wenger, Moraud et al. 2016, Wagner, Mignardot et al. 2018), cortical stimulation (Martinez 2022), and even functional electrical stimulation of leg muscles (Donaldson, Perkins et al. 2000). In the cat, ICMS applied to the motor cortex triggers flexion movements (Bretzner and Drew 2005, Fortier-Lebel, Nakajima et al. 2021), which naturally occur at the beginning of the swing phase during locomotion. Here, we showed that the highest kinematic enhancement and largest reduction in foot drop deficit were observed when the stimulation was delivered in synchrony with the contralateral hindlimb lift. Consistent with previous studies (Bretzner and Drew 2005, Bonizzato and Martinez 2021, Fortier-Lebel, Nakajima et al. 2021), ICMS only affected gait trajectories of the contralateral hindlimb, and this effect was independent of the stimulation timing.

Stimulation burst duration played a smaller, but consistent role, in the control of the foot trajectory. As observed in other studies, short trains of cortical stimulation tend to produce simple motor responses (Brown and Martinez 2018, Brown and Martinez 2021), whereas long trains of stimulation may recruit functionally complex motor responses (Donoghue and Wise 1982, Graziano, Aflalo et al. 2005, Brown and Teskey 2014, Massai, Bonizzato et al. 2021, Brown, Mitra et al. 2022). We tested different durations of stimulation trains, including short (50ms) and longer stimuli (100, 150, 200ms). We found that 100ms trains enhanced natural movement, while limiting the total injected stimulation charge.

Before and after SCI, stimulation amplitude optimization achieved a high-fidelity proportional enhancement of kinematic parameters. The flexion velocity and step height were linearly correlated with increased stimulation amplitude and dragging was linearly reduced after SCI. The proportional stimulation effects were consistent with previous results in rats, both when quantifying cortical population engagement in spontaneous gait modulation (Bonizzato, Pidpruzhnykova et al. 2018) and when applying cortical neurostimulation (Bonizzato and Martinez 2021).

In intact cats, different stimulation sites evoked diverse motor synergies, which could selectively influence leg trajectories during locomotion. After SCI, every site produced a stereotyped modulation of swing, consisting of a flexion movement. Across all animals, this pragmatic vertical motor program proved to be the most resilient against SCI-induced loss of control. This result suggested that fine control of varied motor programs requires a large amount of resources in terms of spared connections. In addition, these data revealed that the multiple neurophysiological processes underpinning spontaneous recovery of gait modulation are focused onto a single compensatory strategy that enhances this retrieved motor program over time. However, the emergence of alternate movement programs cannot be excluded when considering a wider spectrum of less severe SCI.

Phase-coherent neuromodulation can be achieved with a variety of synchronization methods, including real-time kinematic tracking (Wenger, Moraud et al. 2016) and data from tilt sensors or accelerometers (Dai, Stein et al. 1996, Weber, Stein et al. 2004). In this study, we used EMG pattern recognition to decode the extensor muscle activity preceding foot-strike, which was found to be as accurate as video tracking (Wenger, Moraud et al. 2016), with the advantage of being portable (e.g., outside a laboratory setting). With this method, we attained 83% and 88% precision during uni-cortical stimulation in intact and SCI cats, respectively, as well as 97% and 98% precision with bi-cortical stimulation in intact and SCI cats, respectively. The superior pattern recognition performance with bi-cortical stimulation is explained by the regularization effects that phase-coherent cortical stimulation achieves during gait, which is associated with more symmetrical and organized stepping. Indeed, phase-coherent cortical stimulation reduced gait variability by increasing predictability of the lift movement (Bonizzato and Martinez 2021). Bi-cortical neuromodulation applies this regularization effect to both hindlimbs, further benefiting the decoding precision.

Lesion profile analysis revealed that a variable proportion of fibers from cortical and brainstem centers remained intact after SCI. These bridges of intact fibers are essential to convey descending motor commands (generated in the motor cortex) to the lumbar spinal circuits that generate locomotor rhythms (Martinez and Rossignol 2011). We found that the recovery of volitional walking assessed during the obstacle avoidance task paralleled the return of foot clearance. Although spinal plasticity is essential for the recovery of locomotor patterns (Martinez, Delivet-Mongrain et al. 2011, Martinez, Delivet-Mongrain et al. 2012, Martinez, Delivet-Mongrain et al. 2013), the return of volitional walking involves reorganization of the circuits that propagate descending drive generated in the motor cortex (Brown and Martinez 2019). After SCI, cortical (Brown and Martinez 2018, Bonizzato and Martinez 2021, Brown and Martinez 2021), brainstem (Filli, Engmann et al. 2014, Asboth, Friedli et al. 2018), supralesional (Bareyre, Kerschensteiner et al. 2004, Filli and Schwab 2015), and sublesional spinal circuits (Gossard, Delivet-Mongrain et al. 2015) undergo extensive rewiring that are synchronous with recovery.

After SCI, we found that the recovery of locomotor control is time-locked with the return of cortically evoked locomotor responses. As soon as the cats recovered stepping abilities, cortical stimulation evoked hindlimb movements. Since all cats displayed various levels of corticospinal sparing, residual corticospinal fibers may promote transmission of descending drive to spinal circuits. However, we demonstrated that this direct connectivity is not necessary for conveying cortical stimulation in rats with SCI that disrupted all corticospinal fibers (Bonizzato and Martinez 2021). We and others (Asboth, Friedli et al. 2018) have suggested that the upregulation of indirect cortico-reticulospinal pathways, which remain partially spared in our cats, may provide a neural substrate for relaying cortical drive. Other pathways may also be involved. For example, the rubrospinal tract is known to share functional properties with the corticospinal tract during both regulation of fine contralateral forelimb movements (Alstermark, Lundberg et al. 1987, Pettersson, Blagovechtchenski et al. 2000) and modifications of contralateral leg trajectories (Lavoie and Drew 2002). Given the specific effects of cortical stimulation in modulating contralateral movements in real-time, it is unlikely that interhemispheric communication significantly contributed to mediating cortical stimulation effect over locomotor output (Brus-Ramer, Carmel et al. 2009). Direct neuronal activations likely involve localized networks, since current propagation of ICMS depends on its intensity (Stoney, Thompson et al. 1968). Maximal stimuli amplitudes were almost always within the range of 100-300 µA, and never exceeded the 500 µA threshold, which primarily corresponds to sub-millimeter excitation propagation, based on the Stoney equation. Additional mechanistic studies are needed to unveil the pathways through which ICMS exerts its effects over lumbar spinal circuits.

The efficacy of cortical neuroprosthesis in piloting locomotor movements and immediately alleviating bilateral locomotor deficits is here demonstrated in a clinically relevant model of contusive SCI. This research promotes the use of cortical neuromodulation as a movement assistance tool for motor rehabilitation. Similar to our animal model of contusive SCI, individuals with SCI exhibit various lesion and recovery profiles (Frigon 2015), requiring personalized interventions. Our work has shown that cortical neuromodulation exerted a high level of controllability over leg movements and specifically targeted each leg, which allowed for the restoration of a symmetrical gait pattern. Cortical neuroprosthetic approaches may efficiently serve individuals with incomplete SCI or subcortical stroke, who display various degrees of motor deficits, including foot drop and an asymmetrical gait pattern (Mignardot, Le Goff et al. 2017). In the context of rehabilitation, the recovery of voluntary function may be accelerated and enhanced by neuromodulation approaches that repeatedly select, control, and train complex whole-limb lifting movements. Even though implantation and explantation of intracortical probes such as Utah arrays in animals and humans cause minor tissue damage, apart from reactive gliosis (Bullard, Hutchison et al. 2020), the invasive nature of cortical probes remains a hurdle and a potential limitation of our approach. New electrode designs, such as intravascular electrodes (Oxley, Yoo et al. 2021) may mitigate these problems. In addition, non-surgical approaches, including transcranial magnetic stimulation (Benito, Kumru et al. 2012, Kumru, Murillo et al. 2016, Raithatha, Carrico et al. 2016) have been explored in experimental clinical protocols (Dixon, Ibrahim et al. 2016, Jo and Perez 2020). If results are consolidated, then similar protocols may be expanded for broader clinical use. Further experiments are required to establish the trade-off between interface invasiveness and stimulation efficacy. However, it is worth considering that this technique is not more invasive than deep brain stimulation, which has been available for decades and is considered safe for alleviating symptoms of Parkinson’s disease (Stefani, Lozano et al., Liu, Zhang et al.). Moreover, deep brain stimulation is currently being tested in humans with SCI (clinicaltrials.gov ID NCT03053791). As for these approaches, biocompatibility studies are necessary to assess inflammatory responses induced by both our neural interface and chronic delivery of current (Rajan, Boback et al. 2015). In this study, after cortical implantation, we found no motor deficits nor any side effects such as pain.

A major advantage of our animal model is that spinal contusion better reflects the mechanism of injury in humans, as opposed to spinal sections. However, this animal model also has several limitations. First, re-expression of locomotion after severe SCI in cats is more complete compared to humans (Barbeau and Rossignol 1987, Nadeau, Jacquemin et al. 2010, Harnie, Doelman et al. 2019). Spontaneous locomotor recovery in both humans and cats is mediated by several non-exclusive mechanisms, however spinal locomotor circuits may be more autonomous and plastic in the cat (Barbeau and Rossignol 1987, Martinez, Delivet-Mongrain et al. 2011, Martinez, Delivet-Mongrain et al. 2012, Martinez, Delivet-Mongrain et al. 2012, Martinez, Delivet-Mongrain et al. 2013, Harnie, Doelman et al. 2019), since re-expression of spinal locomotion in humans also requires enabling factors (Remy-Neris, Barbeau et al. 1999, Angeli, Edgerton et al. 2014, Rowald, Komi et al. 2022). Nevertheless, the greater reliance of humans on corticospinal input for locomotion (Field-Fote, Yang et al. 2017) only increases the relevance of our supra-lesional stimulation strategy to restore locomotion following an incomplete SCI. Second, cats exhibit significantly less upper motoneuron signs following SCI, notably less spasticity. Despite a few studies, which have shown that repetitive transcranial magnetic stimulation may decrease spasticity in individuals with incomplete SCI (Kumru, Murillo et al. 2010, Benito, Kumru et al. 2012), the underlying neurophysiological mechanisms contributing to this process are still not understood. Consequently, a comprehensive examination of the impact of cortical stimulation on spinal plasticity and excitability is necessary. Lastly, this study only focused on immediate modulation of locomotion, without assessing the long-term effects of cortical stimulation on locomotor recovery. While supported by rat lesion models focused on elucidating the mechanisms of cortical stimulation (Carmel, Berrol et al. 2010, Carmel, Kimura et al. 2014, Bonizzato and Martinez 2021), the long-term effects of bi-cortical neuromodulation on recovery from clinically relevant contusive SCI will require additional studies.

## CRediT authorship contribution statement

Conceptualization: MB, MM; Methodology: MD, MB, HDM, NFL, MM; Investigation: MD, MB, HDM, NFL, MM; Visualization: MD, MB, HDM, MM; Funding acquisition: MM; Project administration: MM; Supervision: MM; Writing – original draft: MD; Writing – review & editing: MD, MB, HDM, NFL, MM.

## Data availability

The data generated throughout this study are available in tabular format as supplementary material. Simple MATLAB code was used to draw figures from these data, and it is available from the authors upon reasonable request. Any additional information required to reanalyze the data reported in this paper is available from the lead contact, Marina Martinez.

## Acknowledgements

The authors would like to thank Gaëlle Forest St-Onge, Alexandre Sheasby, Rania Lejri, David Bergeron, and Anne-Catherine Chouinard for their participation in data processing; Marjolaine Homier, Stéphane Ménard, Raphaël Santamaria, and the staff at the Division des Animaleries for supporting our animal care.

