## Supplementary figures for "Uncovering and exploiting the return of voluntary motor programs after paralysis using a bi-cortical neuroprosthesis"

### Supporting information

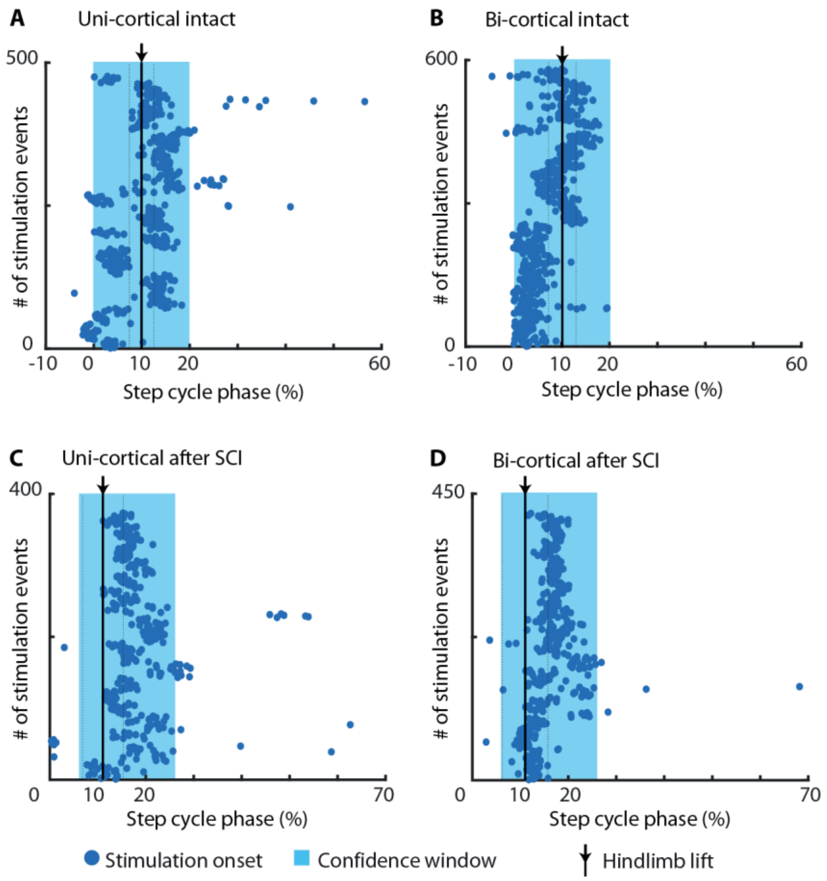


**Fig. S1. Precision of EMG activity pattern recognition to enable timely delivery of stimulation in the intact state and after SCI.** Phase of the step cycle in which stimulation onset occurred when **(A)** uni-cortical and **(B)** bi-cortical stimulation was delivered during locomotion in the intact state. Phase of the step cycle in which stimulation onset occurred when **(C)** uni-cortical and **(D)** bi-cortical stimulation was delivered during locomotion following bilateral spinal cord contusion. Precision takes into consideration the proportion of stimulation events whose onsets falls inside the expected time window. This window corresponds to 20% of the step cycle in which stimulation results in the highest step height in the intact state, and highest reduction of dragging deficits after SCI.

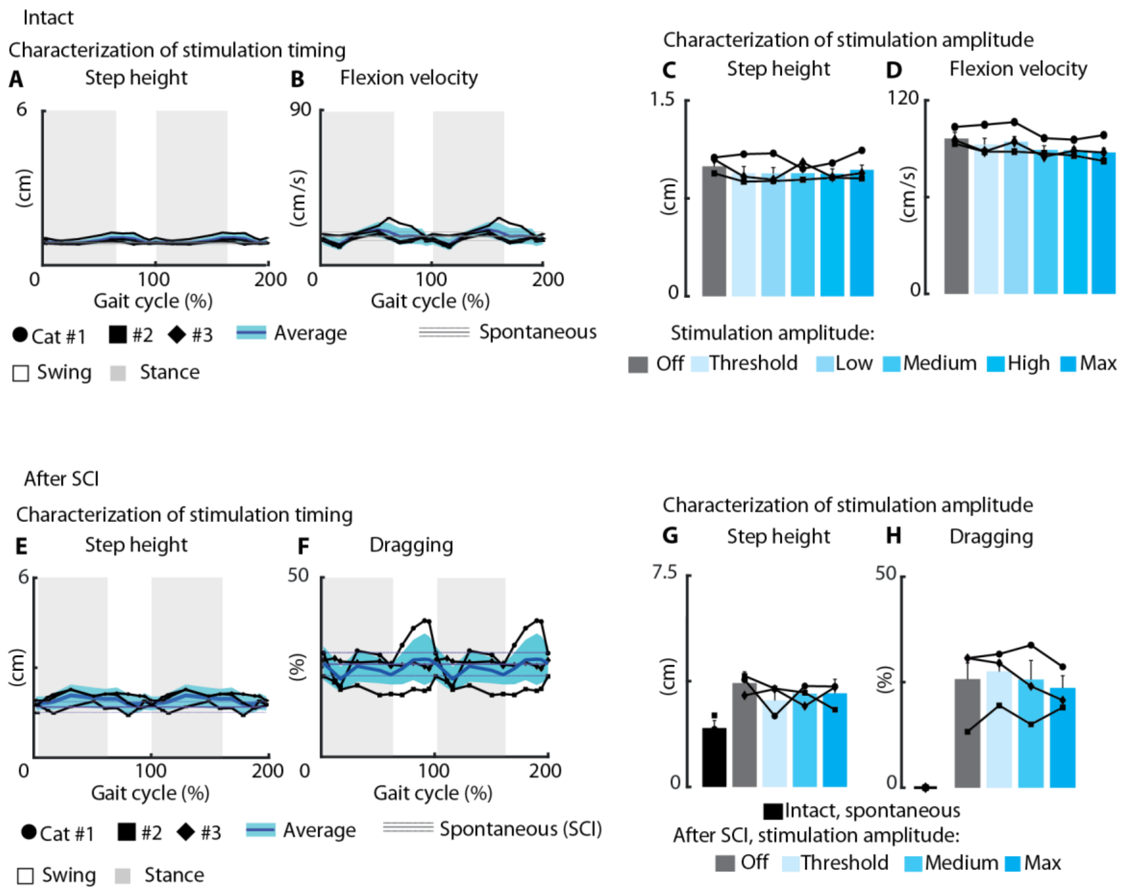


**Fig. S2. Uni-cortical stimulation did not modulate the kinematics of the ipsilateral hindlimb in the intact state and after contusive SCI. (A-D)** In intact cats, neither the timing nor the amplitude of ICMS significantly modulated step height and flexion velocity of the ipsilateral hindlimb. **(E-H)** Consistently with the results observed in the intact state, neither the timing nor the amplitude of ICMS significantly modulated step height and dragging of the ipsilateral hindlimb after SCI.

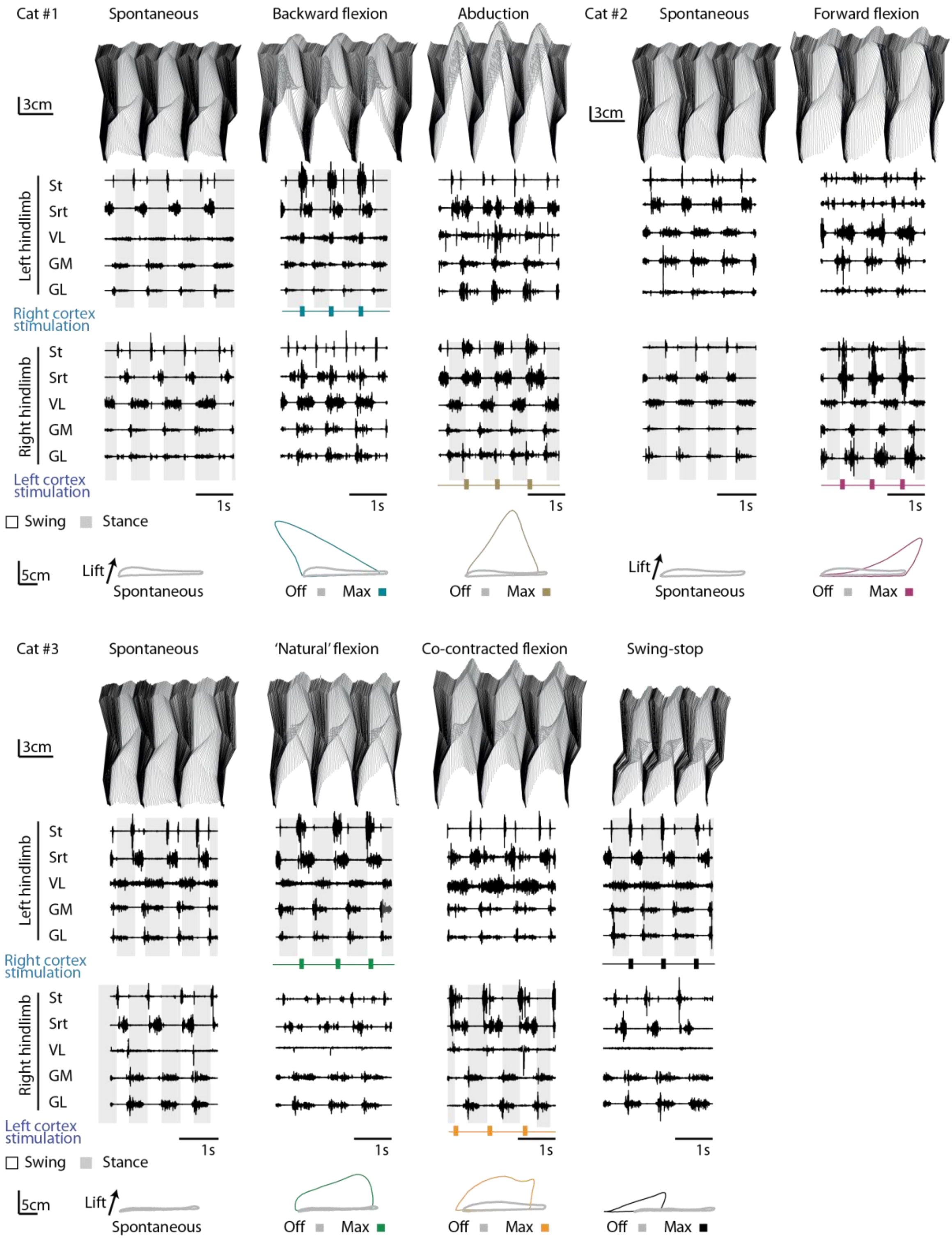


**Fig. S3. Uni-cortical stimulation triggered a variety of motor programs in intact cats.** Each sub-panel presents a stick diagram, muscle activity and limb trajectory during spontaneous walking (no stimulation), or during phase-coherent ICMS.

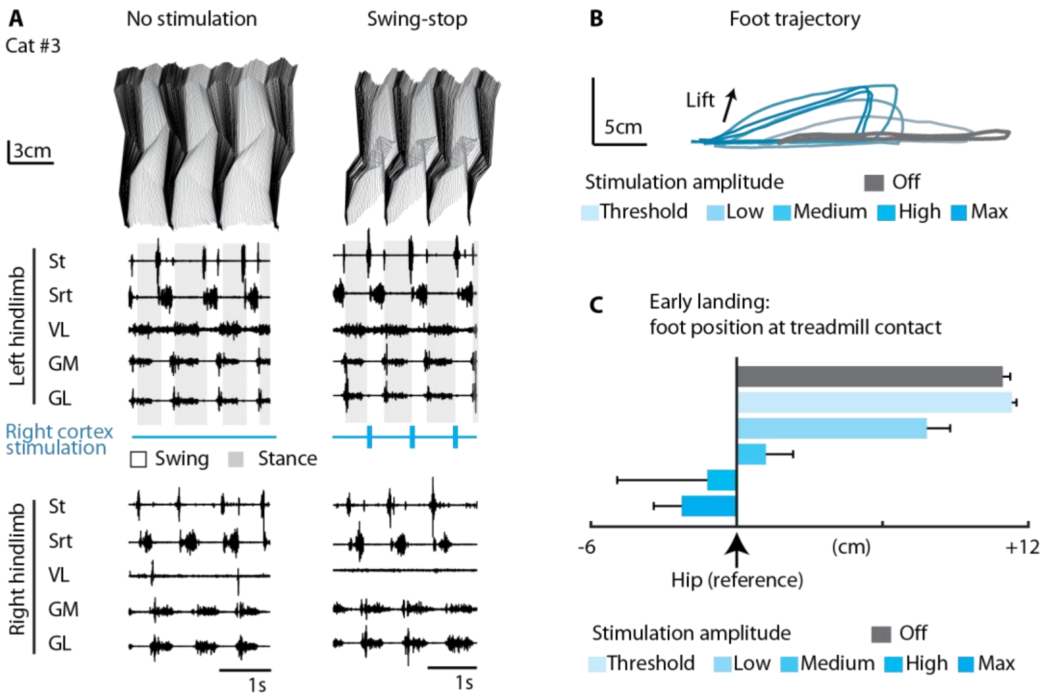


**Fig. S4. Case study of an intracortical stimulation site that produced a swing-stop movement. (A)** Stick figure of the left hindlimb during spontaneous walking and when stimulation was delivered to the right cortex. **(B)** The stimulation amplitude delivered at this site modulated foot trajectories by shortening the swing. **(C)** With increasing stimulation amplitudes, the foot landed on the treadmill proportionally earlier.

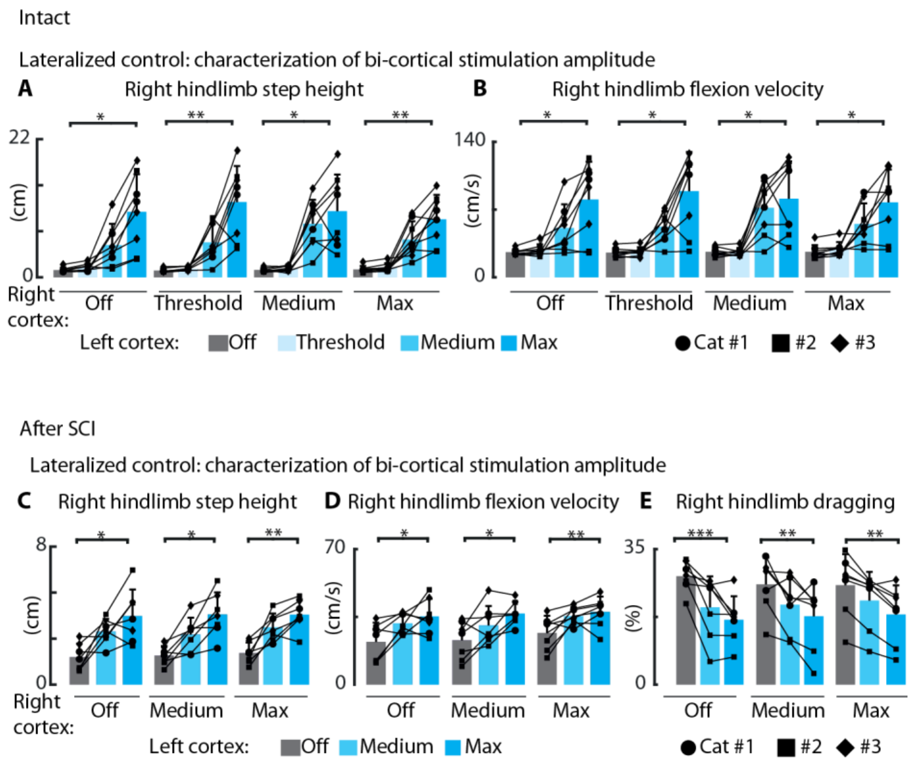


**Fig. S5. Bi-cortical stimulation modulated bilateral hindlimb movements in intact and SCI cats and reduced dragging deficits after SCI.** This Figure complements Figure 5C-D and Figure 6D-E, presenting the results obtained for the right hindlimb in lieu of the left one. In intact cats, during bi-cortical stimulation, right hindlimb **(A)** step height and **(B)** Flexion velocity linearly modulated with increasing ICSM amplitude applied to the left cortex, independently of the stimulation amplitude delivered to the right cortex. Consistently with the results obtained in the intact state, right hindlimb **(C)** step height, **(D)** Flexion velocity and **e** reduction of dragging linearly modulated with increasing ICMS amplitude applied to the left cortex after SCI, independently of the stimulation amplitude delivered to the right cortex.

**Movie S1:** Movement variety elicited by cortical stimulation at the intact state and after SCI.

**Movie S2:** Bi-cortical stimulation modulates locomotion at the intact state.

**Movie S3**: Bi-cortical stimulation alleviates bilateral locomotor deficits after SCI.
